# Antibiotics can be used to contain drug-resistant bacteria by maintaining sufficiently large sensitive populations

**DOI:** 10.1101/638924

**Authors:** Elsa Hansen, Jason Karslake, Robert J. Woods, Andrew F. Read, Kevin B. Wood

## Abstract

Standard infectious disease practice calls for aggressive drug treatment that rapidly eliminates the pathogen population before resistance can emerge. When resistance is absent, this *elimination strategy* can lead to complete cure. However, it also removes drug-sensitive cells as quickly as possible, removing competitive barriers that may slow the growth of resistant cells. In contrast to the elimination strategy, the *containment strategy* aims to maintain the maximum tolerable number of pathogens, exploiting competitive suppression to achieve chronic control. Here we combine *in vitro* experiments in computer-controlled bioreactors with mathematical modeling to investigate whether containment strategies can delay failure of antibiotic treatment regimens. To do so, we measured the “escape time” required for drug-resistant *E. coli* populations to eclipse a threshold density maintained by adaptive antibiotic dosing. Populations containing only resistant cells rapidly escape containment, but we found that matched populations with the maximum possible number of sensitive cells could be contained for significantly longer. The increase in escape time occurs only when the threshold density–the acceptable bacterial burden–is sufficiently high, an effect that mathematical models attribute to increased competition. The findings provide decisive experimental confirmation that maintaining the maximum number of sensitive cells can be used to contain resistance when the size of the population is sufficiently large.

## Introduction

The ability to successfully treat infectious disease is often undermined by the emergence of drug resistance [1–6]. When resistance poses a major threat to the quality and duration of a patient’s life, as it can in chronic infections, the goal of treatment is to restore patient health while delaying treatment failure for as long as possible. To do so, standard practice calls for aggressive drug treatment to rapidly remove the drug-sensitive pathogen population to prevent resistance-conferring mutations [7–17]. This *elimination strategy* also dominates cancer therapy, where treatment is often aimed at achieving rapid and dramatic reductions in tumor burden [18–29]. Elimination is the best way to stop *de novo* resistance, and if resistance is absent, it can lead to a complete cure. On the other hand, if resistance is already present, or it is not possible to treat sufficiently, then aggressive treatment may actually promote the emergence of resistance [30–39]. Indeed, it is clear from the substantial number of treatment failures associated with drug resistance in both cancers and infectious diseases that an aggressive approach does not always work [1, 4–6].

If aggressive treatment cannot prevent the emergence of resistance, an alternative approach is to use competition between drug-sensitive and drug-resistant cells to slow this emergence. There is ample evidence that competition between sensitive and resistant cells can be intense [40–43] and may be over limiting resources like glucose or target cells [44–47]. Competition can also be immune-mediated or occur via direct interference (e.g. bacteriocins) [40, 48–51]. There are numerous theoretical studies [49, 52–60] showing that sensitive cells can competitively suppress resistant cells, and this suppression has been observed experimentally in parasites and cancer [33, 36–38, 56, 61, 62]. Since sensitive cells can both generate *de novo* resistance and also competitively suppress existing resistant mutants, making good treatment decisions requires understanding the relative importance of these opposing effects (Figure 1).

**Fig 1.**
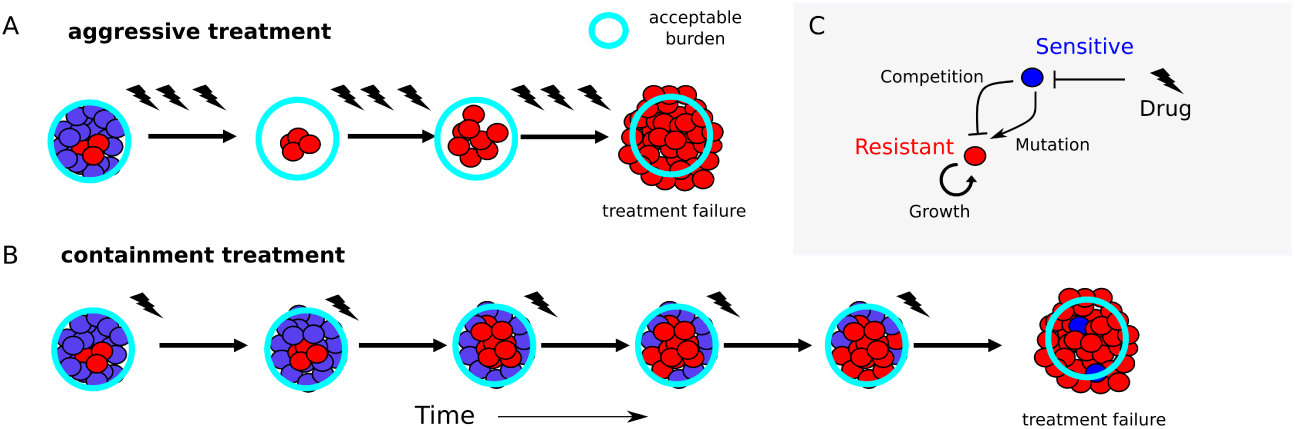
Containment strategies may leverage competition to extend time below failure threshold. A. Aggressive treatment uses high drug concentrations (lightning flashes), which eliminates sensitive cells (blue) but may fail when resistant cells (red) emerge and the population exceeds the failure threshold (“acceptable burden”, light blue circle). B. Containment strategies attempt to maintain the population just below the failure threshold, leveraging competition between sensitive (blue) and emergent resistant (red) cells to potentially prolong time to failure. C. Schematic of potential feedback between growth processes in mixed populations. Drug (lightning flash) inhibits sensitive cells (blue), which in turn inhibit resistant cells (red) through competition but may also contribute to the resistant population via mutation.

To address this, recent theoretical work compares two extreme treatment strategies: a mutation minimizing strategy (what we call elimination) and a competition maximizing strategy (what we call containment) [59]. The elimination strategy removes sensitive cells as fast as possible, minimizing the risk of mutation but also removing competitive barriers that may slow the growth of existing resistant cells [49] (Figure 1A). Containment, on the other hand, maintains as many sensitive pathogens as is clinically acceptable (i.e., a pathogen density that is deemed to be safe and below which treatment is not necessary), using drug treatment only to alleviate symptoms. Containment maintains the total pathogen density at this acceptable burden (Figure 1B). This maximizes competitive suppression but leaves sensitive pathogens which can generate resistance. In a sense, all possible drug regimens must lie somewhere on the spectrum between elimination (mutation minimization) and containment (sensitive maximization). Cases where containment better delays resistance emergence represent situations where standard practice can potentially be improved.

Theory predicts that neither strategy is always best; there are situations where containment will better control resistance emergence and others where it will make a bad prognosis worse [59]. The latter case arises when the benefit of maintaining sensitive pathogens (competitive suppression of resistance) is outweighed by the rate at which sensitive pathogens become resistant by mutation (or horizontal gene transfer). Mathematical modeling suggests that across a wide variety of diseases and settings, the same fundamental principles determine when containment is better than elimination. First, there has to be sufficient competition (the clinically acceptable burden high enough) and second, the growth of the resistant population has to be driven primarily by the replication of resistant cells, not by mutational inputs (resistant population large enough).

Here we experimentally test the hypothesis that containment can delay the emergence of resistant bacteria. We combine simple mathematical models with *in vitro* experiments in computer-controlled bioreactors, where “treatment” failure is defined by bacterial populations eclipsing a threshold density (the acceptable burden). The experimental design directly tests mutation minimization against competition maximization and demonstrates that containment type strategies can increase escape times when the acceptable burden is sufficiently high. Our experiments provide a clear test of the idea that competition can be harnessed to control resistance. Although there is empirical evidence for competitive suppression in parasites and cancer [33, 36–38, 61, 62], we believe this is the first explicit demonstration of competitive suppression of resistance to antibiotic treatment. We are also unaware of any direct tests of containment, the competition-maximizing strategy, in any pathogen. The findings are particularly striking because they occur in well mixed populations with a continual renewal of resources and using an acceptable burden well below the natural carrying capacity - all conditions not typically associated with strong competition.

## Results

The aim of this study is to provide “proof-of-concept” that, under appropriate conditions, containment can better manage antibiotic resistance than elimination. This is done by developing an experimental system where mutation minimization can be directly compared to competition maximization.

### Model System

We grew bacterial populations in well mixed bioreactors where environmental conditions, including drug concentration and nutrient levels, can be modulated using a series of computer-controlled peristaltic pumps (see, for example, [63, 64]). Population density is measured using light scattering (optical density, OD), and drug concentration can be adjusted in real time in response to population dynamics or predetermined protocols (Figure S1). This setup allows us to implement mutation minimizing and competition maximizing strategies on bacterial populations with different drug sensitivities.

To obtain bacterial populations with different drug sensitivities, we began with *E. coli* strains REL606 and REL607, which are well-characterized ancestral strains used in the celebrated long term evolution experiment in *E. coli* [65]. These strains differ by a single point mutation in *araA* which serves as a neutral marker for competition experiments; REL 606 (REL 607) appears red (pink) when grown on tetrazolium arabinose (TA) plates. To generate a “drug-resistant” strain, we used laboratory evolution to isolate a mutant of the REL606 strain that was resistant to doxycycline, a frequently used protein synthesis inhibitor (Methods). We then quantified the responses of the drug-sensitive (REL607) and drug-resistant (REL606-derived mutant) cells to doxycycline by measuring real-time per capita growth rate for isogenic populations of each strain exposed to different concentrations of drug (Figure 2). Briefly, growth rate was estimated using influx rate of media required to maintain populations at a constant density (Methods). The resistant isolate exhibits both increased resistance to doxycycline (increased half maximal inhibitory concentration) as well as decreased growth in the absence of drug (Figure 2). We note that in the experiments that follow, drug concentrations are sufficiently high that resistant cells generally have a selective advantage over sensitive cells, despite this fitness cost.

**Fig 2.**
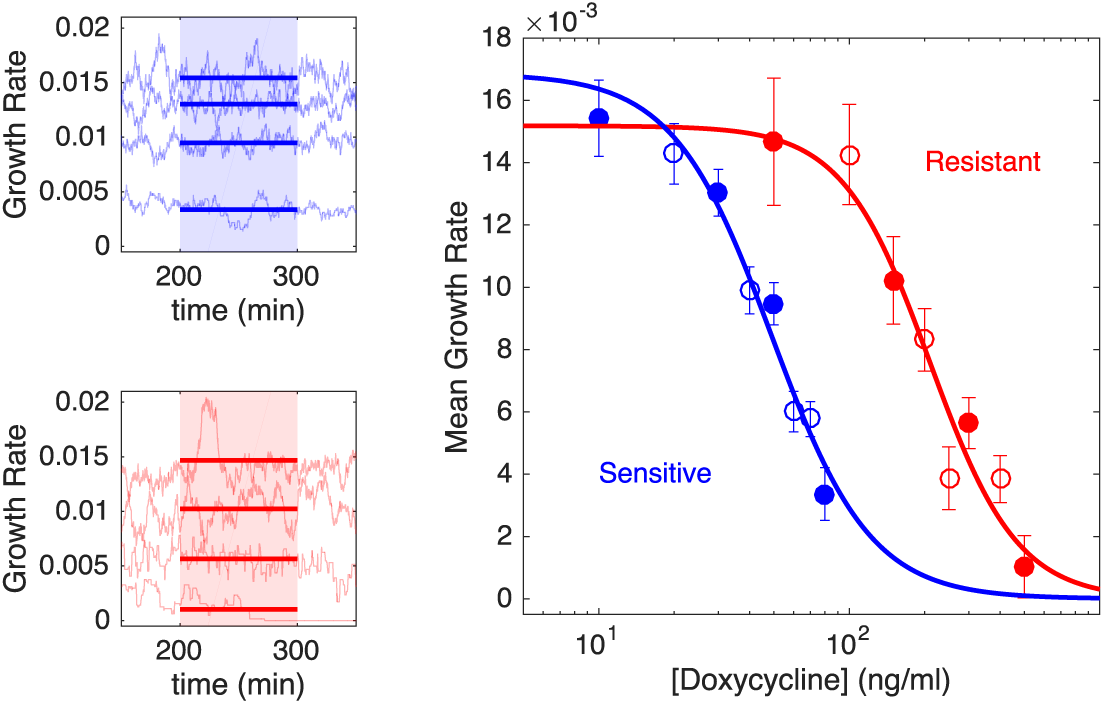
Resistant cells exhibit increased resistance to doxycycline and small fitness cost. Left panels: per capita growth rate in bioreactors for ancestral (sensitive, blue) and resistant (red) populations exposed to increasing concentrations of doxycycline (top to bottom in each panel). Real time per capita growth rate (light blue or red curves) is estimated from flow rates required to maintain constant cell density at each drug concentration (Methods). Mean growth rate (thick solid lines) is estimated between 200-300 minutes post drug addition (shaded regions), when the system has reached steady state. Doxycycline concentrations are 10, 30, 50, and 80 ng/mL (top panel, top to bottom) and 50, 150, 300, and 500 ng/mL (bottom panel, top to bottom). Right panel: dose response curve for sensitive (blue) and resistant (red) populations. Circles are time averaged growth rates between 200-300 minutes post drug addition (shaded regions in A), with error bars ± one standard deviation over the measured interval; filled circles correspond to the specific examples shown in left panels. Drug-free growth rates are 0.018 ± 0.001 min^*−*1^ (sensitive) and 0.015 ± 0.002 min^*−*1^ (resistant), indicating that resistant cells exhibit a fitness cost. Solid lines, fit to Hill-like dose response function *r* = *r*_0_(1 + (*D/h*)^*k*^)^*−*1^, with *r* the growth rate, *D* the drug concentration, *r*_0_ the growth in the absence of drug, *h* the half-maximal inhibitory concentration (IC_50_), and *k* the Hill coefficient. Half-maximal inhibitory concentrations are estimated to be *h* = 49 ng/mL (sensitive cells) and *h* = 210 ng/mL (resistant cells).

### Experimental Design

With this model system we are able to implement “mutation minimization” and “competition maximization” strategies. An idealized version of mutation minimization is achieved by monitoring a bacterial population without any sensitive cells (Figure 3A: resistant-only). Because there are no sensitive cells in the initial population, the potential for mutational input (i.e. sensitive cells mutating to become resistant) is minimized. Competition maximization is achieved by implementing a drug-dosing protocol that maintains a mixed population with the largest possible sensitive density (Figure 3A: Mixed).

**Fig 3.**
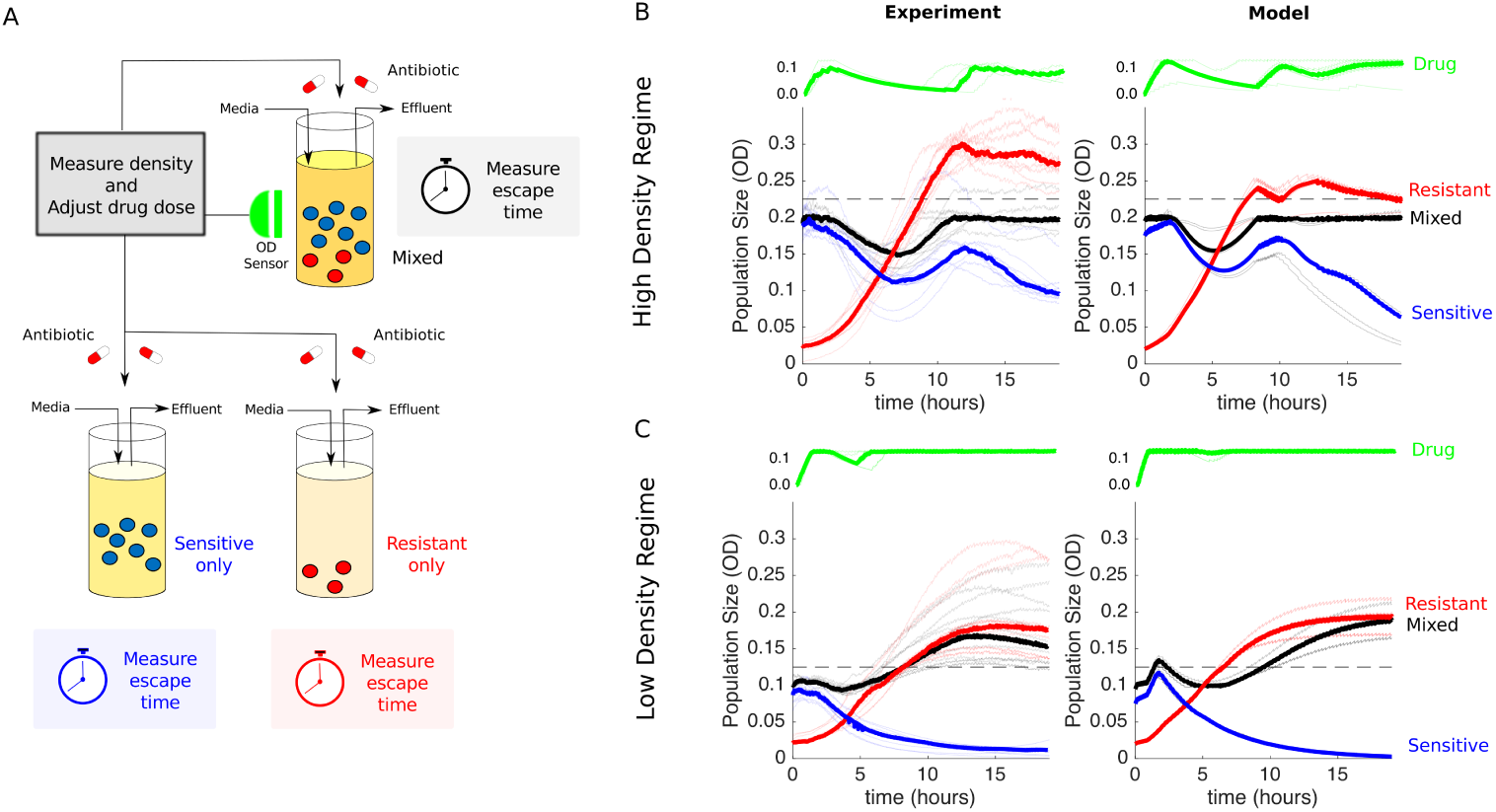
Experimental design and dynamics for comparison of “mutation minimization” and “competition maximization”. A. Schematic of experiment. Three different populations (sensitive-only, resistant-only, and mixed) were exposed to identical antibiotic treatments in separate bioreactors. The media in each bioreactor was also refreshed at a constant rate of 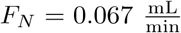. The drug treatment was determined in real time by measuring the density (OD) of the mixed population and adjusting drug influx to maintain a constant density (P_max_) while minimizing drug used (Methods). While the dynamics of the mixed population fully determine the temporal profile of the drug dosing, all 3 populations then receive identical treatments. In the high density experiment, mixed populations started at an OD of P_max_ = 0.2, with a 90-10 ratio of sensitive to resistant cells. The initial OD of resistant cells is therefore 0.02. Resistant-only populations started from an initial density of 0.02 and contained no sensitive cells, while sensitive-only populations started from an initial density of 0.18 and contained no resistant cells. In the low density experiment, mixed populations started at an OD of P_max_ = 0.1, and the initial OD of resistant cells was unchanged (OD=0.02). Therefore, the starting conditions of the high and low density experiments differ only in the number of sensitive cells. B and C. Experiments (left) and model (right) in high density (P_max_ = 0.2, B) and low density (P_max_ = 0.1, C) regimes. Red curves are resistant-only, blue are sensitive-only, and black are the mixed populations. Lightly shaded curves correspond to individual experiments, dark curves show the median across experiments. Horizontal dashed lines show the treatment failure threshold P_max_ + 0.025, where the 0.025 term allows for small experimental fluctuations without triggering a threshold crossing event.

The experiment is then the following. First, we seed two vials with the same low density of resistant cells. Then, drug sensitive cells are added to the ‘mixed’ vial, to achieve a total cell density equal to a predetermined threshold density – the acceptable burden, *P*_max_. Because high concentrations of drug are expected to completely inhibit growth of sensitive cells and therefore eliminate any potential competition, we designed an adaptive drug dosing protocol intended to maintain the mixed population at a fixed density (P_max_, which we call the acceptable burden) using minimal drug. The dosing protocol uses simple feedback control to adjust the drug concentration in real time in response to changes in population density (Figure 3A, Methods). Finally, as a control, a third vial is seeded with the same density of sensitive bacteria as that added to the mixed vial (Figure 3A: sensitive-only). This control allows for an indirect measure of the effect of sensitive cells in the mixed vial since, in the absence of competition or other intercellular interactions, the dynamics of the mixed population should be a simple sum of the dynamics in the two single-species populations.

The temporal dynamics of the mixed population–but not the other populations– completely determine the drug dosing protocol, but all three populations receive identical drug dosing and therefore experience identical drug concentrations over time. This strategy ensures that any increased resistance suppression in the mixed vial can be attributed to competitive suppression by sensitive cells and not drug inhibition. Because drug concentration in the vials is restricted to a finite range (0-125 ng/mL), populations containing resistant cells cannot be contained indefinitely and will eventually eclipse the target density (P_max_). The time required for this crossover is defined as the escape time, and the goal of the experiment is to compare escape times–which correspond, intuitively, to times of treatment failure–under mutation minimizing and competition maximizing conditions.

There are three possibilities. If mutation dominates, then mutation minimization is expected to be best and the escape time of the resistant-only population should exceed that of the mixed-population. On the other hand, if competition dominates then the mixed population should take the longest to escape. Finally, if the effects of sensitive cells (both mutational input and competitive suppression) are negligible then the escape times of the resistant-only and mixed populations should be similar. To quantitatively guide our experiments and refine this intuition, we developed and parameterized a simple mathematical model for population growth in the bioreactors (see Table 1, Methods and SI for a detailed description of the model). A simple analysis suggests that, for our experimental design, the effect of competition will always dominate over the effect of mutational input (see SI). As a result, we neglect mutation and focus on the role of competition. This allows us to identify two values of acceptable burden (P_max_) which are predicted to produce different results. For high *P*_max_ (OD= 0.2), competition dominates and the mixed vial should have the longest escape time. For low *P*_max_ (OD= 0.1), competition is minimal and the escape times of the resistant-only and mixed vials should be similar.

**Table 1.** Model Parameter Description

| Parameter | Definition | Value |
| --- | --- | --- |
| $V$ | volume of vial | 17 mL |
| $F_D$ | flow rate of drug reservoir | 1 $\frac{\text{mL}}{\text{min}}$ |
| $\chi_D$ | function indicating when drug is being added | 1 when drug is being added<br>0 when drug is not being added |
| $F_N$ | constant background flow rate of nutrients | 0.067 $\frac{\text{mL}}{\text{min}}$ |
| $D_{in}$ | drug concentration in drug reservoir | 500 $\frac{\text{ng}}{\text{mL}}$ |
| $C$ | carrying capacity | 280000000 $\frac{\text{bacteria}}{\text{mL}}$ ($OD = 0.35$) |
| $r_S$ | intrinsic per capita growth rate of drug sensitive strain | 0.0169 $\frac{1}{\text{min}}$ |
| $r_R$ | intrinsic per capita growth rate of drug resistant strain | 0.0152 $\frac{1}{\text{min}}$ |
| $h_S$ | drug concentration where effect of drug is at 50% (assuming $h = 1$) | 49.0639 $\frac{\text{ng}}{\text{mL}}$ |
| $h_R$ | drug concentration where effect of drug is at 50% (assuming $h = 1$) | 209.9995 $\frac{\text{ng}}{\text{mL}}$ |
| $k_S$ | hill function coefficient | 2.2023 |
| $k_R$ | hill function coefficient | 2.4849 |
| $\tau_S$ | time delay for sensitive strain | 79.04 min |
| $\tau_R$ | time delay for resistant strain | 96.72 min |

### Benefit of competition maximization depends on acceptable burden

To test these predictions, we first performed the experiment at the threshold density that the model predicts will lead to competitive suppression (P_max_ = 0.2 (Figure 3B)). Note that this density falls in the range of exponential growth and falls below the stationary phase limit in unperturbed populations (Figure S2). To account for batch effects and day-to-day experimental fluctuations, we repeated the experiment multiple times across different days, using different media and drug preparations. Unsurprisingly, the experiments confirm that sensitive-only populations are significantly inhibited under this treatment protocol and never reach the containment threshold; in fact, the overall density decreases slowly over time due to a combination of strong drug inhibition and effluent flow (Figure 3B, blue curves). By contrast, the resistant-only population grows steadily and eclipses the threshold in 6-9 hours (Figure 3B, red curves). Remarkably, however, the mixed population (black curves) is contained below threshold–in almost all cases–for the entire length of the experiment, which spans more than 18 hours. At the end of the experiment, we plated representative examples of resistant-only and mixed populations (Figure S3), which confirmed that the mixed vial was predominantly resistant at the end of the experiment. In addition, these sensitive and resistant isolates exhibited similar dose-response curves to the original sensitive and resistant strains (Figure S5). Matched drug-free controls indicate that containment in the mixed vial is due to drug, not artifacts from media inflow or outflow (Figure S3). The experiments also show remarkable agreement with the model (with no adjustable parameters; compare left and right panels in Figure 3B).

If competition were driving the increased escape time, one would expect the effect to be reduced as the threshold density (P_max_) is decreased. To test this hypothesis, we repeated the experiments at P_max_ = 0.1 (Figure 3C). As before, the sensitive-only population is strongly inhibited by the drug and decreases in size over time (blue). Also as before, the resistant-only population (red) escapes the containment threshold, typically between 5-8 hours (faster than in the high P_max_ experiment due to the lower threshold). In contrast to the previous experiment, however, the mixed population also escapes the containment threshold, and furthermore, it does so on similar timescales as the resistant-only population. Again, the agreement between model and experiment is quite good, though the model does predict a slightly longer escape time in the mixed population. The small discrepancy between the model and the experiment suggests that competition may decrease even more rapidly as density is lowered than the model assumes.

To quantify these results, we calculated the time to escape for each experiment. We defined time to escape for a particular experiment as the first time at which the growth curve (OD) exceeded the threshold density P_max_ by at least 0.025 OD units (note that the 0.025 was chosen to allow for noise fluctuations in the OD time series without triggering a threshold crossing event). For low values of acceptable burden (P_max_), the escape times for resistant-only and mixed populations are nearly identical (Figure 4, left). By contrast, at higher values of P_max_, the escape time is dramatically increased in the mixed population relative to the resistant-only population (Figure 4, right), even though both receive identical drug treatment and start with identically sized resistant populations. Importantly, our experiments suggest that sensitive cells are beneficial at high values of P_max_ and have little effect at low P_max_, consistent with the assumption that mutation-driven costs of sensitive cells in our system are negligible and that the acceptable burden must be high enough to derive benefit from competitive suppression.

**Fig 4.**
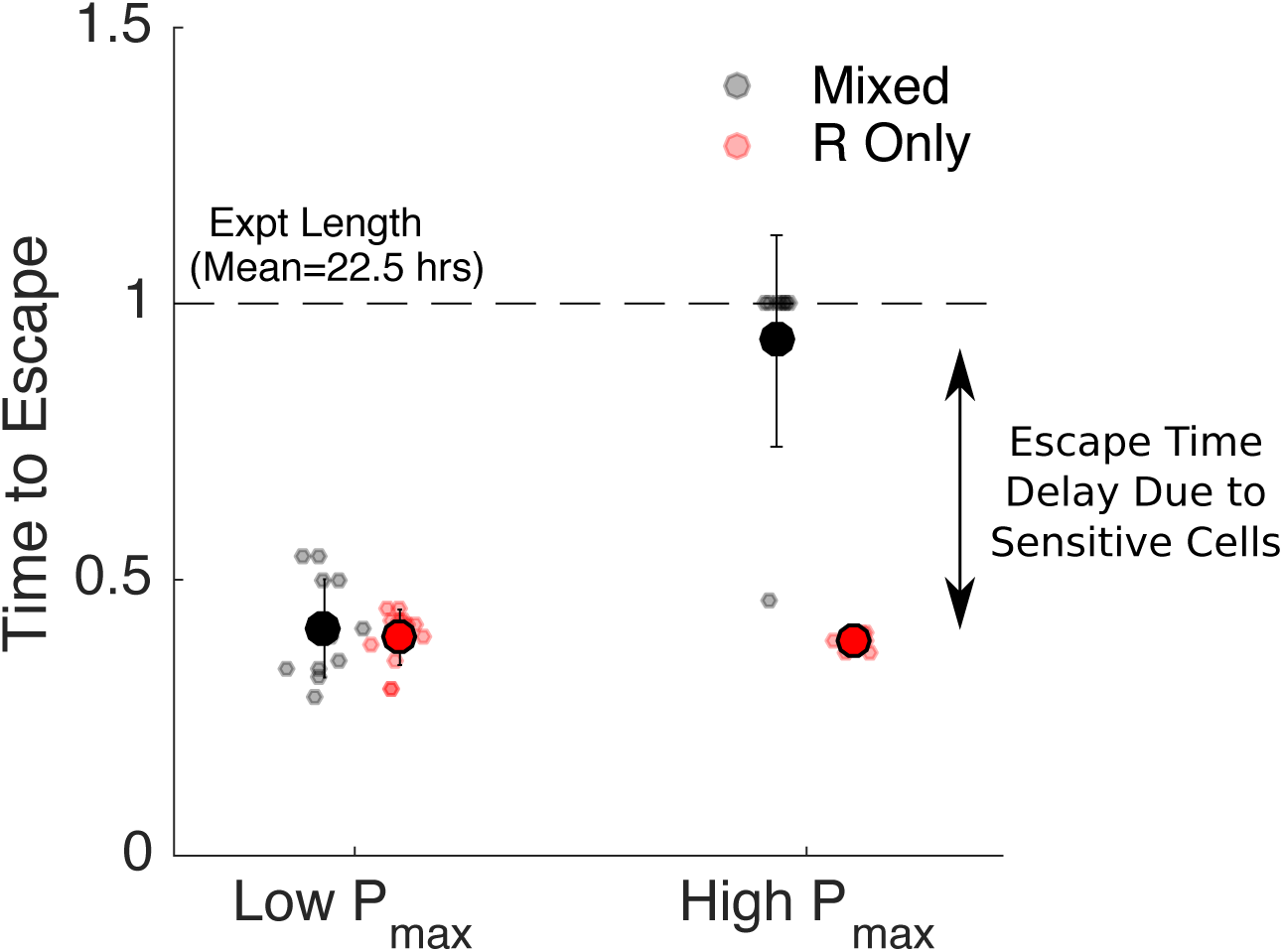
Increased escape time under “competition maximization” requires a high threshold density. Time to escape for populations maintained at low (P_max_ = 0.1, left) and high (P_max_=0.2, right) threshold densities (“acceptable burdens”). Small circles: escape times for individual experiments in mixed (black) or resistant-only (red) populations. Large circles: mean escape time across experiments, with error bars corresponding to *±* one standard deviation. Time to escape is defined as the time at which the population exceeds the threshold OD of P_max_ + 0.025, where the 0.025 is padding provided to account for noise fluctuations. Time to escape is normalized by the total length of the experiment (mean length 22.5 hours). Note that in the high P_max_ case (right), the mixed population (black) reached the threshold density during the course of the experiment in only one case, so the escape times are set to 1 in all other cases.

## Discussion

In this work, we provide direct experimental evidence that the presence of drug-sensitive cells can lead to improved antibiotic-driven control of bacterial populations *in vitro*. Specifically, we show that a “competition maximizing” strategy can contain mixed populations of sensitive and resistant cells below a threshold density for significantly longer than matched populations containing only resistant cells. The increase in escape time occurs only when the threshold density is sufficiently high that competition is significant. The findings are particularly remarkable given that experiments are performed in well mixed bioreactors with continuous resource renewal, and even the highest density thresholds occur in the exponential growth regime for unperturbed populations. The surprisingly strong effect of competition under these conditions suggests that similar approaches may yield even more dramatic results in natural environments, where spatial heterogeneity and limited diffusion may enhance competition [66–68].

Notably, our experiments did not uncover scenarios where sensitive cells may actually be detrimental and accelerate resistance emergence. Theory suggests that these scenarios do indeed exist [59], but because of the typical mutation rates observed in bacteria, they cannot be reliably produced with our experimental system (see SI for extended discussion).

It is important to keep in mind several technical limitations of our study. First, we measured population density using light scattering (OD), which is a widely used experimental surrogate for microbial population size but is sensitive to changes in cell shape [69]. Because we use protein synthesis inhibitors primarily at sub-MIC concentrations, we do not anticipate significant artifacts from this limitation, though it may pose challenges when trying to extend these results to drugs such as fluoroquinolones, which are known to induce filamentation [70, 71]. In addition, in the absence of cell lysis, OD can not distinguish between dead and living cells. However, our experiments include a slow background flow that adds fresh media and removes waste, leading to a clear distinction between non-growing and growing populations. Under these conditions, fully inhibited (or dead) populations would experience a decrease in OD over time, while populations maintained at a constant density are required to divide at an effective rate equal to this background refresh rate.

Most importantly, our results are based entirely on *in vitro* experiments, which allow for precise environmental control and quantitative measurements but clearly lack important complexities of realistic *in vivo* and clinical scenarios. Developing drug protocols for clinical use is an extremely challenging problem. Our goal was not to design clinically realistic containment strategies, but instead to provide direct experimental evidence that protocols aimed at maximally maintaining sensitive cells can delay resistance emergence in a tractable setting. The drug dosing protocol applied here attempts to supply the minimum possible amount of drug to control the population. However, the results show that it still reduces the size of the sensitive population, suggesting that the protocol could be improved. As a result, our experimental results are likely underestimating the potential benefit of maximizing the sensitive population. The fact that we are still able to detect a benefit to maintaining (a non maximal) sensitive population indicates that there is room for implementing these types of strategies in the non-idealized setting of real-life. In fact, adaptive strategies designed to leverage competition between drug-sensitive and drug-resistant cancer cells have been successful both in mouse models [56] and in clinical trials [60]. Our results suggest that these successes may be further improved by designing “competition maximizing” strategies.

A basic science understanding of containment has the potential for real world impact, as containment strategies are being tested on people now. If our hypothesis is confirmed, there are definable situations where existing strategies could be improved by maximizing competition as opposed to simply using “less aggressive” adaptive approaches. We hope these experiments will motivate continued experimental, theoretical, and perhaps even clinical investigations, particularly in situations where the primary threat to the well-being of the patient is the *de novo* emergence of drug resistance (e.g. [60, 72]).

## Methods

### Bacterial Strains, Media, and Growth Conditions

Experiments were performed with *Escherichia coli* strains REL 606 and 607 [65]. A resistant strain (“resistant mutant”) was isolated from lab-evolved populations of REL606 undergoing daily dilutions (200X) into fresh media with increasing doxycycline (Research Products International) concentrations for 3 days. A single resistant isolate was used for all experiments. Stock solutions were frozen at −80C in 30 percent glycerol and streaked onto fresh agar plates (Davis Minimal Media (Sigma) with 2000 µg/ml glucose) as needed. Overnight cultures of resistant and sensitive cells for each experiment were grown from single colonies and then incubated in sterile Davis Minimal Media with 1000 µg/ml glucose liquid media overnight at 30C while rotating at 240 rpm. All bioreactor experiments were performed in a temperature controlled warm room at 30C.

### Continuous Culture Bioreactors

Experiments were performed in custom-built, computer-controlled bioreactors as described in [64], which are based, in part, on similar designs from [63, 73]. Briefly, constant volume bacterial cultures (17 mL) are grown in glass vials with customized Teflon tops that allow inflow and outflow of fluid via silicone tubing. Flow is managed by a series of computer-controlled peristaltic pumps—up to 6 per vial—which are connected to media and drug reservoirs and allow for precise control of various environmental conditions. Cell density is monitored by light scattering using infrared LED/Detector pairs on the side of each vial holder. Voltage readings are converted to optical density (OD) using a calibration curve based on separate readings with a table top OD reader. Up to 9 cultures can be grown simultaneously using a series of multi-position magnetic stirrers. The entire system is controlled by custom Matlab software.

### Experimental Mixtures and Set up

Before the experiments begin, vials are seeded with sensitive or resistant strains of *E. coli* and allowed to grow to the desired density in the bioreactor vials. Cells were then mixed (to create the desired population compositions) and diluted as appropriate to achieve the desired starting densities. Each vial is connected to 1) a drug reservoir containing media and doxycycline (500 *µ*g/ml), 2) a drug-free media reservoir that provides constant renewal of media, 3) an effluent waste reservoir. Flow from reservoir 1 (drug reservoir) is determined in real-time according to a simple feedback algorithm intended to maintain cells at a constant target density with minimal drug. Flow to/from reservoirs 2 and 3 provides a slow renewal of media and nutrients while maintaining a constant culture volume in each vial.

### Drug Dosing Protocols

To determine the appropriate antibiotic dosing strategy the computer records the optical density in each vial every 3 seconds. Every 3 minutes, the computer computes:(i) the average optical density *OD*_*avg*_ in the mixed vial over the last 30 seconds and (ii) the current drug concentration in the vial. If *OD*_*avg*_ is greater than *P*_*max*_, the desired containment density, and the current drug concentration is less than 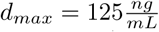 then drug and media will be added to the vial for 21 seconds at a flow rate of 1 mL/min. In a typical experiment, this control algorithm is applied to one of the mixed populations to determine, in real time, the drug dosing protocol (i.e. influx of drug solution over time). The exact same drug dosing protocol is then simultaneously applied to all experimental populations (mixed, resistant-only, sensitive-only, Figure 3). In parallel, an identical dosing protocol is applied to a series of control populations, but in these populations, the drug solution is replaced by drug-free media (Figure S3).

### Mathematical Model

The mathematical model used in the simulations is

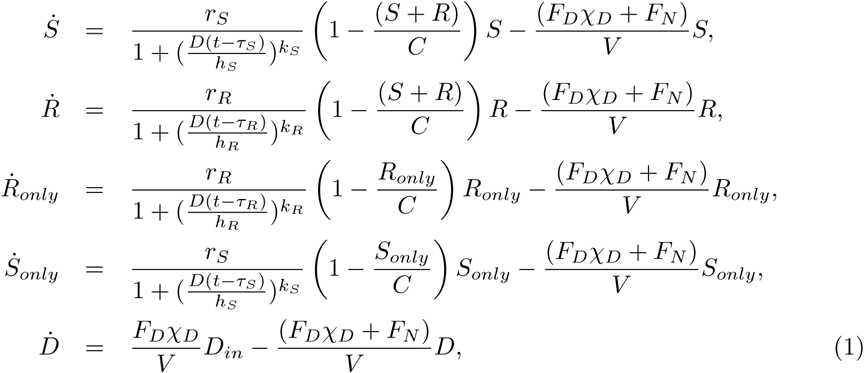

where *S* and *R* are the drug-sensitive and drug-resistant densities in mixed vial, *R*_*only*_ is the bacterial density in the vial that contains only drug-resistant bacteria, *S*_*only*_ is the bacterial density in the vial that contains only drug-sensitive bacteria and *D* is the drug concentration in the vials,. The initial conditions for the simulations (and experiments) are given in Table 2. The effect of drug on growth rate is modeled as a hill function with parameters *r*_*S*_, *k*_*S*_ and *h*_*S*_ for the sensitive strain and parameters *r*_*R*_, *k*_*R*_ and *h*_*R*_ for the resistant strain. There is also a time-delay associated with the effect of drug (denoted by *τ*_*S*_ for the sensitive strain and *τ*_*R*_ for the resistant strain). Competition in the model is captured by using a logistic growth term with carrying capacity *C*. It is assumed that the sensitive and resistant strains have similar carrying capacities. Finally, the bioreactor has a continual efflux to maintain constant volume. The rate of this outflow is the sum of the constant background nutrient flow *F*_*N*_ and any additional outflow required to compensate for the inflow of drug which enters at a rate *F*_*D*_*χ*_*D*_. *F*_*D*_ is a constant rate and *χ*_*D*_ is an indicator function which is 1 when drug is being added to the vials and 0 when it is not. In the simulations, the decision of when to add drug is based on the same control algorithm that was used in the actual experiment (see Methods: Drug Dosing Protocols). Since Model (1) describes the rate of change of bacterial density, the total efflux (*F*_*N*_ + *F*_*D*_*χ*_*D*_) is divided by the volume of the vials *V*. The drug concentration in the vials is determined by the rate of drug flow into the vials (*F*_*D*_*χ*_*D*_*D*_*in*_, where *D*_*in*_ is the concentration of drug in the reservoir) and the rate of efflux out of the vials (*F*_*D*_*χ*_*D*_ + *F*_*N*_). The values of *D*_*in*_, *V*, *F*_*D*_ and *F*_*N*_ were chosen to match the associated values in the experimental system, all other parameters in the model where fit using independent experimental data (see SI for details) and are given in Table 1.

**Table 2.**
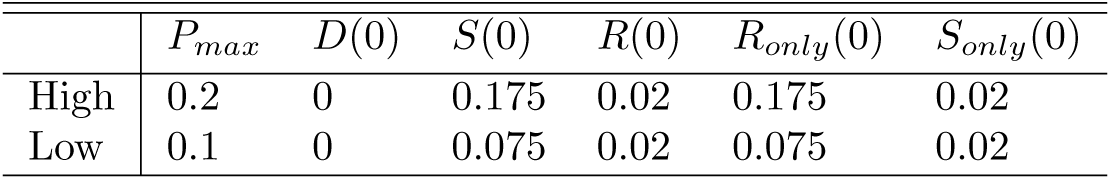
Initial Optical Densities for Simulations and Experiments

## Acknowledgements

This work is supported by the National Science Foundation (NSF No. 1553028 to KW), the National Institutes of Health (NIH No. 1R35GM124875-01 to KW; NIH No. R01 GM089932 to AFR; NIH K08 AI119182 to RW), and the Eberly Family (to AFR). The funders had no role in study design, data collection and analysis, decision to publish, or preparation of the manuscript. The format of the preprint is based on a modified PLoS latex template provided on Overleaf.com.

## Supplemental Information

### S1 Drug Free Control Populations

In addition to the experimental populations grown in each bioreactor experiment (described in Figure 3A), we also measured growth in 3 control populations grown in neighboring bioreactor vials. One vial contained the resistant strain (starting at the same initial density as in the resistant-only experimental population) but experienced no influx or outflow of media and received no drug. These unperturbed growth curves are shown in Figure S2. A second control vial contained the resistant strain (again starting at the same initial density as in the resistant-only experimental population) and received inflow and outflow identical to that received in the experimental populations. However, the drug inflow solution was replaced by drug-free media. A third vial was identical to the second, but also contained sensitive cells and therefore served as a drug-free matched control of the mixed experimental population. The latter two controls eclipse the density threshold in each experiment (Figure S3), indicating that the adaptive therapies leading to containment inhibit growth via drug rather than via rapid influx and outflow.

### S2 Parameter Estimates

The resistant and sensitive strains were characterized using independent experiments that measured growth at different drug concentrations. Using a method detailed in [64], the CCD was configured to hold bacterial populations growing in different concentrations of drug at a constant density. The pump rate was fixed at 1 mL/min and then the pumps were turned on and off in order to maintain a constant OD. Growth rates were then determined by the average amount of time the pumps were on - pumps on more often indicates faster growth rates. If *r*_*S*_ (*D*) and *r*_*R*_(*D*) are the growth rates of the sensitive and resistant strains at drug concentration *D* then:

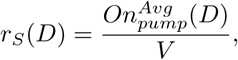

and

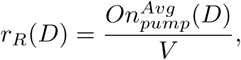

where *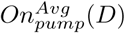* is the average amount of time the pumps are on for drug concentration *D* and *V* is the volume of the vial.

Using these estimated growth rates, we then fit a function (similar to a hill function) to characterize how each strain responds to different concentrations of drug. Specifically,

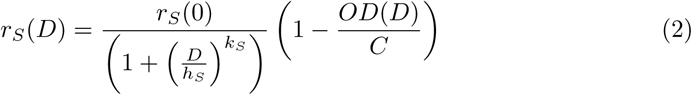

and

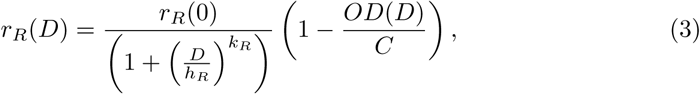

where *OD*(*D*) is the the constant *OD* that the the population was held at when exposed to drug concentration *D*. Although different experiments were held at slightly different constant densities, *OD*(*D*) was always very close to 0.02. In Equations (2) and (3), the term 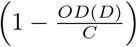 is included to account for the fact that there may still be some competition at this low density.

In general, 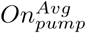 was higher at the beginning of an experiment and then decreased to a constant value, indicating that there is a time delay for drug to take effect. To account for this we used data between minutes 200 and 300 of the experiment to fit the above equations. By this time, the drug had reached its full effect and 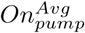 was essentially constant.

We also estimated the time delay associated with drug effect and included this in the main model (Model (1) from main text). The time delay for drug effect was estimated using the following equation:

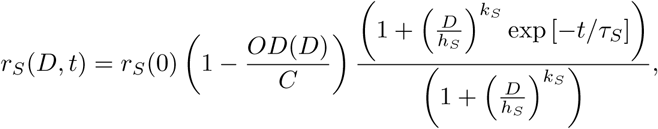

and

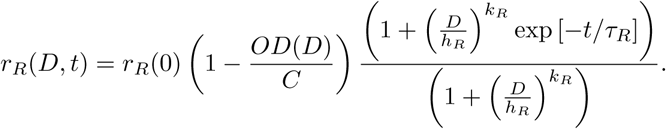

These equations were fit using all the data from an experiment that corresponded to a constant bacterial density. From these fits we obtained a time delay for the effect of drug for the sensitive (*τ*_*S*_) and the resistant (*τ*_*R*_) strain.

### S3 Considering Mutation

Our experiments were seeded with a resistant OD of 0.02 (approximately 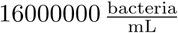). With this choice, the probability of mutation to resistance would have to be at least 0.056, and in practice much higher, before the cost of mutation would outweigh the benefit of competition. Alternatively, if the probability of mutation to resistance was 0.001 then the relative proportion of resistance at the beginning of the experiment would have to be less than 1 in 1000 (a situation that cannot be reliably created in our system) before there was even a possibility that maintaining a sensitive population would be detrimental. This means that within the operating regime of our experimental system and for the specific characteristics of the sensitive and resistant strains used, maintaining a sensitive population will never be detrimental. These calculations are described in more detail below.

Using a slightly modified version of our mathematical model of the bioreactor (Model 1 in main text), we can estimate what the probability of mutation would need to be before the presence of sensitive cells was detrimental. The dynamics of the resistant density in the mixed vial is given by:

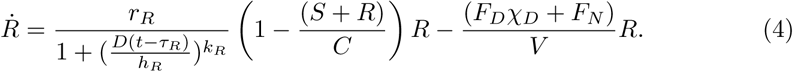

If the probability of mutation to resistance is ϵ, then adding mutational input to Equation (4) resiults in:

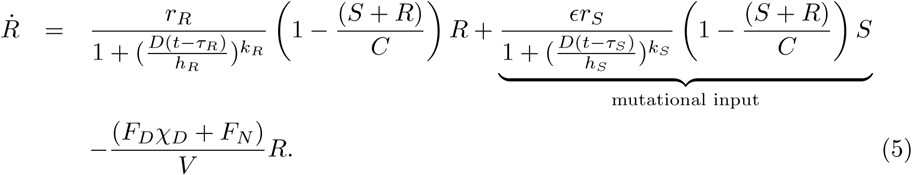

Equation (5) can be rewritten to isolate the effect that sensitives have on the resistant population:

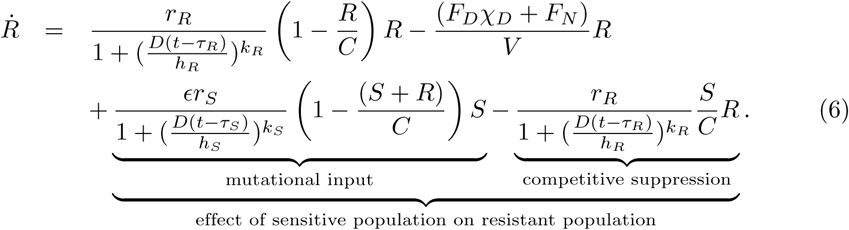

Therefore, the benefit of competitive suppression will exceed the cost of mutation whenever

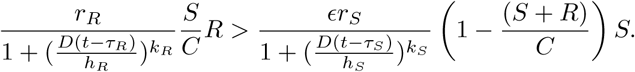

We can use the above relation to obtain a lower bound for how much mutation there must be before there is any risk of the sensitive population being detrimental. Specifically,

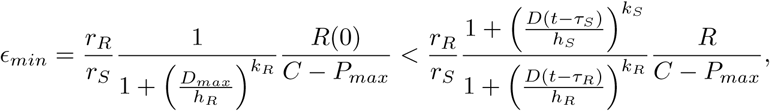

where 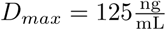 is the maximum drug concentration allowed in the vials and *R*(0) is the starting resistant density. Using the parameter values for our model this gives *ϵ*_*min*_ = 0.056 when *P*_*max*_ = 0.1 and *ϵ*_*min*_ = 0.094 when *P*_*max*_ = 0.2.

Note that *ϵ*_*min*_ is a lower bound for how high the probability of mutation must be before sensitive cells are detrimental. In practice, *ϵ* could be higher than this and the net effect of the sensitive population (over the entire course of the infection) could still be to delay resistance emergence.

Alternatively, if we know the probability of mutation to resistance then we can compute an upper bound for the starting resistant density:

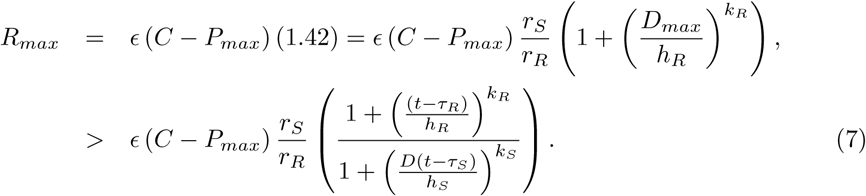

Therefore, if the probability of mutation to resistance is 0.001 then the starting OD for the resistant strain would have to be less than 0.0002 (for *P*_*max*_ = 0.2) and less than 0.00035 (for *P*_*max*_ = 0.1) before there is any risk of mutational costs exceeding competitive benefits.

**Fig S1.**
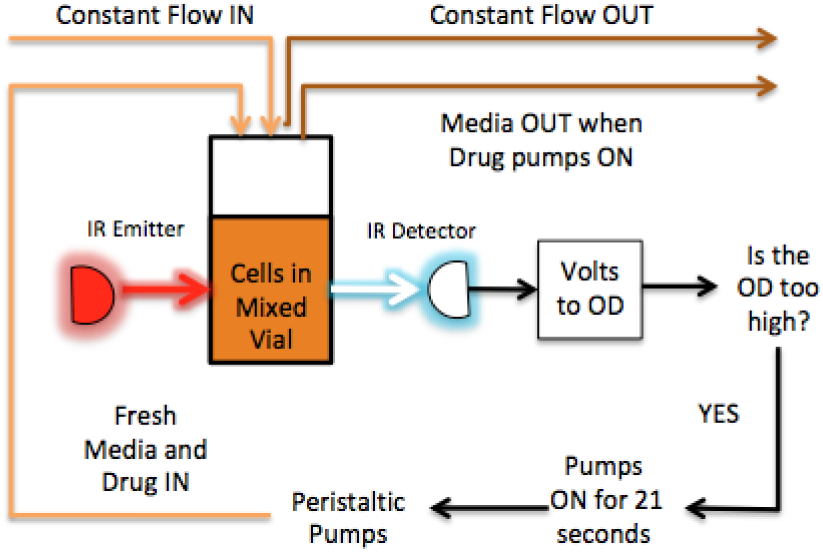
Computer controlled bioreactors. Constant volume bacterial cultures (17 mL) are grown in glass vials with customized Teflon tops that allow inflow and outflow of fluid via silicone tubing. Flow is managed by a series of computer-controlled peristaltic pumps which are connected to media and drug reservoirs. Cell density is monitored by light scattering using infrared LED/Detector pairs on the side of each vial holder. Voltage readings are converted to optical density (OD) using a calibration curve based on separate readings with a table top OD reader. Up to 9 cultures can be grown simultaneously using a series of multi-position magnetic stirrers. The entire system is controlled by custom Matlab software. Flow chart (above) depicts adaptive drug therapy (lower branches) intended to maintain constant OD by adding drug in response to changes in cell density.

**Fig S2.**
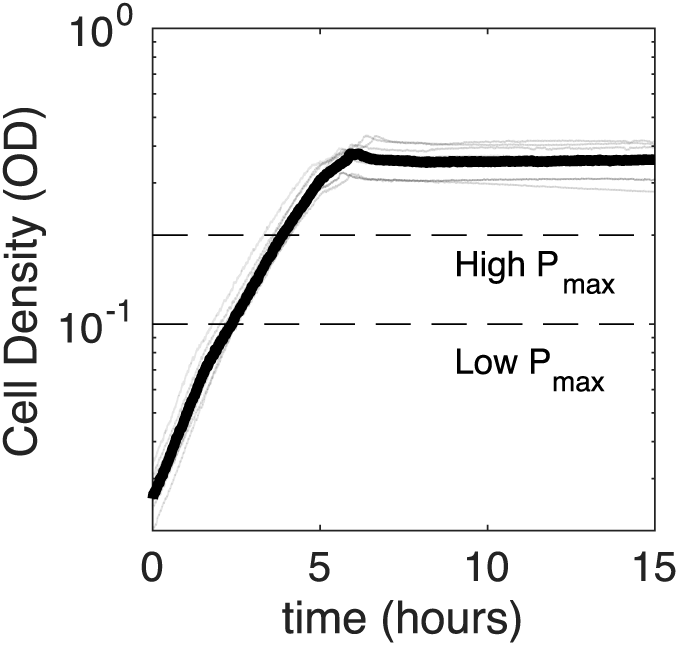
Growth of resistant cells in unperturbed bioreactors. Cell density (OD) over time for REL607-derived resistant strains in bioreactors without influx or outflow of media. Transparent black lines correspond do growth curves performed in parallel with each bioreactor experiment. Thick black curve is the median over replicates. Dashed lines indicate threshold densities used in experiments (P_max_ = 0.2 and P_max_ = 0.1).

**Fig S3.**
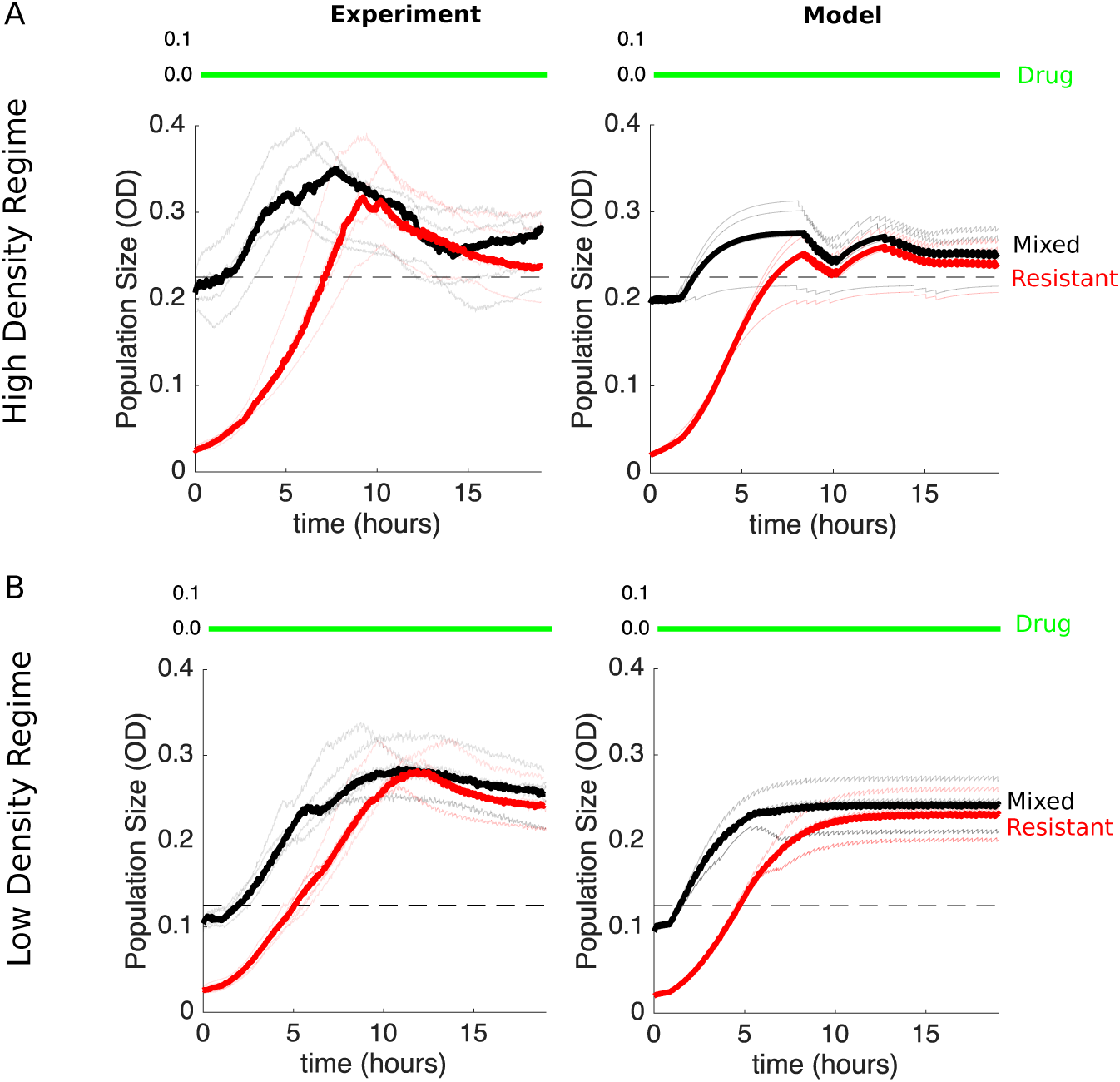
Matched drug-free control populations are not contained by adaptive dosing protocol. Conditions are identical to those in Figure 3B and Figure 3C except that all populations receive drug-free media rather than drug solution media as part of the adaptive dosing protocol.

**Fig S4.**
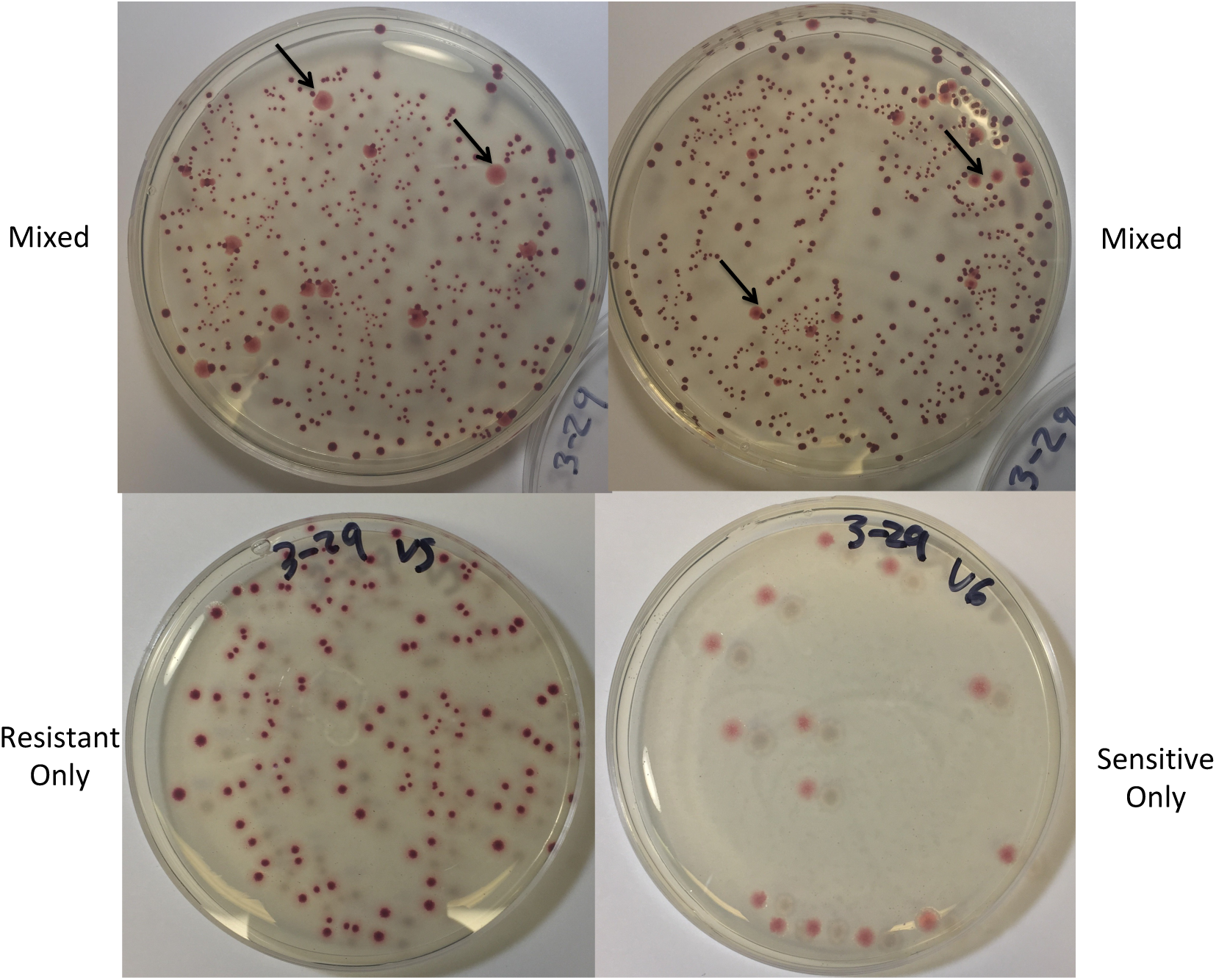
Mixed populations contain primarily resistant cells at final time point of escape time experiment. The REL606-derived resistant strain appears red and the sensitive REL607 strain appears pink when grown on tetrazolium arabinose (TA) plates. Upper panels: Samples from two mixed vials taken at the end of a high density escape time experiment (as in Figure 3B). Arrows indicate sensitive colonies. Bottom row: Samples from the end of a high density escape time experiment for a vial seeded with only resistant bacteria (left) and a vial seeded with only sensitive bacteria (right).

**Fig S5.**
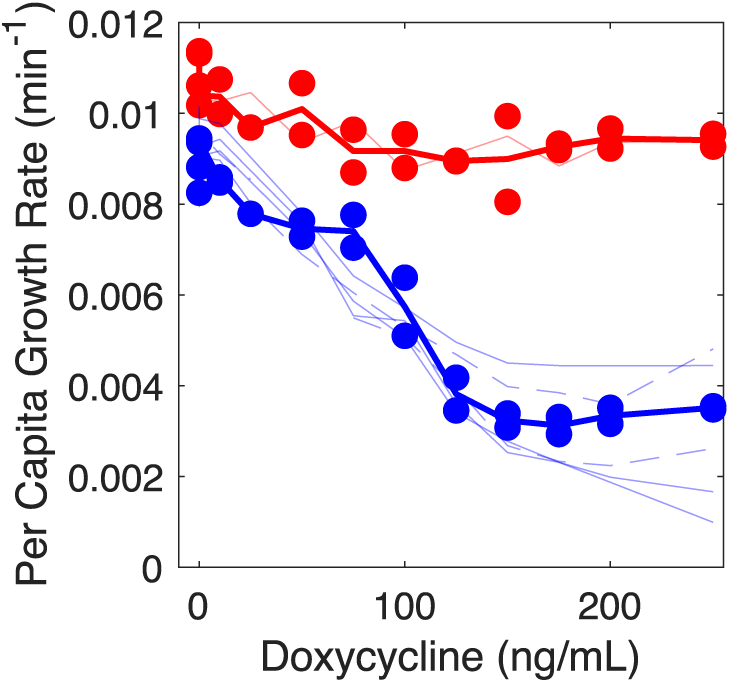
Sensitive and resistant isolates show similar dose-response curves before and after experiment. Dose response curves measured in 96-well microplates for sensitive (REL607; blue circles) and REL606-derived resistant strains (red circles). Dark line, mean across replicates. Thin (transparent) curves correspond to colonies isolated from population mixture at the end of an escape time experiment. Red curve, resistant isolate (appears red on plate); Blue curves, sensitive isolates (appear pink on plate).

