## Supplemental Text for "Antibiotics can be used to contain drug-resistant bacteria by maintaining sufficiently large sensitive populations"

### Supplemental Information

#### S1 Drug Free Control Populations

In addition to the experimental populations grown in each bioreactor experiment (described in Figure 3A), we also measured growth in 3 control populations grown in neighboring bioreactor vials. One vial contained the resistant strain (starting at the same initial density as in the resistant-only experimental population) but experienced no influx or outflow of media and received no drug. These unperturbed growth curves are shown in Figure S2. A second control vial contained the resistant strain (again starting at the same initial density as in the resistant-only experimental population) and received inflow and outflow identical to that received in the experimental populations. However, the drug inflow solution was replaced by drug-free media. A third vial was identical to the second, but also contained sensitive cells and therefore served as a drug-free matched control of the mixed experimental population. The latter two controls eclipse the density threshold in each experiment (Figure S3), indicating that the adaptive therapies leading to containment inhibit growth via drug rather than via rapid influx and outflow.

#### S2 Parameter Estimates

The resistant and sensitive strains were characterized using independent experiments that measured growth at different drug concentrations. Using a method detailed in [71], the CCD was configured to hold bacterial populations growing in different concentrations of drug at a constant density. The pump rate was fixed at 1 mL/min and then the pumps were turned on and off in order to maintain a constant OD. Growth rates were then determined by the average amount of time the pumps were on - pumps on more often indicates faster growth rates. If $r_S(D)$ and $r_R(D)$ are the growth rates of the sensitive and resistant strains at drug concentration $D$ then:

$$r_S(D) = \frac{On_{pump}^{Avg}(D)}{V},$$

and

$$r_R(D) = \frac{On_{pump}^{Avg}(D)}{V},$$

where $On_{pump}^{Avg}(D)$ is the average amount of time the pumps are on for drug concentration $D$ and $V$ is the volume of the vial.

Using these estimated growth rates, we then fit a function (similar to a hill function) to characterize how each strain responds to different concentrations of drug. Specifically,

$$r_S(D) = \frac{r_S(0)}{\left(1 + \left(\frac{D}{h_S}\right)^{k_S}\right)} \left(1 - \frac{OD(D)}{C}\right) \tag{2}$$

and

$$r_R(D) = \frac{r_R(0)}{\left(1 + \left(\frac{D}{h_R}\right)^{k_R}\right)} \left(1 - \frac{OD(D)}{C}\right), \tag{3}$$

where $OD(D)$ is the the constant $OD$ that the the population was held at when exposed to drug concentration $D$. Although different experiments were held at slightly different constant densities, $OD(D)$ was always very close to 0.02. In Equations (2) and (3), the term $\left(1 - \frac{OD(D)}{C}\right)$ is included to account for the fact that there may still be some competition at this low density.

In general, $On_{pump}^{Avg}$ was higher at the beginning of an experiment and then decreased to a constant value, indicating that there is a time delay for drug to take effect. To account for this we used data between minutes 200 and 300 of the experiment to fit the above equations. By this time, the drug had reached its full effect and $On_{pump}^{Avg}$ was essentially constant.

We also estimated the time delay associated with drug effect and included this in the main model (Model (1) from main text). The time delay for drug effect was estimated using the following equation:

$$r_S(D,t) = r_S(0)\left(1 - \frac{OD(D)}{C}\right)\frac{\left(1 + \left(\frac{D}{h_S}\right)^{k_S}\exp\left[-t/\tau_S\right]\right)}{\left(1 + \left(\frac{D}{h_S}\right)^{k_S}\right)},$$

and

$$r_R(D,t) = r_R(0)\left(1 - \frac{OD(D)}{C}\right)\frac{\left(1 + \left(\frac{D}{h_R}\right)^{k_R}\exp\left[-t/\tau_R\right]\right)}{\left(1 + \left(\frac{D}{h_R}\right)^{k_R}\right)}.$$

These equations were fit using all the data from an experiment that corresponded to a constant bacterial density. From these fits we obtained a time delay for the effect of drug for the sensitive ($\tau_S$) and the resistant ($\tau_R$) strain.

#### S3 Considering Mutation

Our experiments were seeded with a resistant OD of 0.02 (approximately 16000000 $\frac{\text{bacteria}}{\text{mL}}$). With this choice, the probability of mutation to resistance would have to be at least 0.056, and in practice much higher, before the cost of mutation would outweigh the benefit of competition. Alternatively, if the probability of mutation to resistance was 0.001 then the relative proportion of resistance at the beginning of the experiment would have to be less than 1 in 1000 (a situation that cannot be reliably created in our system) before there was even a possibility that maintaining a sensitive population would be detrimental. This means that within the operating regime of our experimental system and for the specific characteristics of the sensitive and resistant strains used, maintaining a sensitive population will never be detrimental. These calculations are described in more detail below.

Using a slightly modified version of our mathematical model of the bioreactor (Model 1 in main text), we can estimate what the probability of mutation would need to be before the presence of sensitive cells was detrimental. The dynamics of the resistant density in the mixed vial is given by:

$$\dot{R} = \frac{r_R}{1 + (\frac{D(t-\tau_R)}{h_R})^{k_R}}\left(1 - \frac{(S+R)}{C}\right)R - \frac{(F_D\chi_D + F_N)}{V}R. \tag{4}$$

If the probability of mutation to resistance is $\epsilon$, then adding mutational input to Equation (4) resiults in:

$$\dot{R} = \frac{r_R}{1 + (\frac{D(t-\tau_R)}{h_R})^{k_R}}\left(1 - \frac{(S+R)}{C}\right)R + \underbrace{\frac{\epsilon r_S}{1 + (\frac{D(t-\tau_S)}{h_S})^{k_S}}\left(1 - \frac{(S+R)}{C}\right)S}_{\text{mutational input}} - \frac{(F_D\chi_D + F_N)}{V}R. \tag{5}$$

Equation (5) can be rewritten to isolate the effect that sensitives have on the resistant population:

$$\begin{aligned}\dot{R} &= \frac{r_R}{1+(\frac{D(t-\tau_R)}{h_R})^{k_R}}\left(1-\frac{R}{C}\right)R - \frac{(F_D\chi_D+F_N)}{V}R \\ &\quad + \underbrace{\underbrace{\frac{\epsilon r_S}{1+(\frac{D(t-\tau_S)}{h_S})^{k_S}}\left(1-\frac{(S+R)}{C}\right)S}_{\text{mutational input}} - \underbrace{\frac{r_R}{1+(\frac{D(t-\tau_R)}{h_R})^{k_R}}\frac{S}{C}R}_{\text{competitive suppression}}}_{\text{effect of sensitive population on resistant population}}. \qquad (6)\end{aligned}$$

Therefore, the benefit of competitive suppression will exceed the cost of mutation whenever

$$\frac{r_R}{1+(\frac{D(t-\tau_R)}{h_R})^{k_R}}\frac{S}{C}R > \frac{\epsilon r_S}{1+(\frac{D(t-\tau_S)}{h_S})^{k_S}}\left(1-\frac{(S+R)}{C}\right)S.$$

We can use the above relation to obtain a lower bound for how much mutation there must be before there is any risk of the sensitive population being detrimental. Specifically,

$$\epsilon_{min} = \frac{r_R}{r_S}\frac{1}{1+\left(\frac{D_{max}}{h_R}\right)^{k_R}}\frac{R(0)}{C-P_{max}} < \frac{r_R}{r_S}\frac{1+\left(\frac{D(t-\tau_S)}{h_S}\right)^{k_S}}{1+\left(\frac{D(t-\tau_R)}{h_R}\right)^{k_R}}\frac{R}{C-P_{max}},$$

where $D_{max} = 125\frac{\text{ng}}{\text{mL}}$ is the maximum drug concentration allowed in the vials and $R(0)$ is the starting resistant density. Using the parameter values for our model this gives $\epsilon_{min} = 0.056$ when $P_{max} = 0.1$ and $\epsilon_{min} = 0.094$ when $P_{max} = 0.2$.

Note that $\epsilon_{min}$ is a lower bound for how high the probability of mutation must be before sensitive cells are detrimental. In practice, $\epsilon$ could be higher than this and the net effect of the sensitive population (over the entire course of the infection) could still be to delay resistance emergence.

Alternatively, if we know the probability of mutation to resistance then we can compute an upper bound for the starting resistant density:

$$\begin{aligned}R_{max} &= \epsilon\,(C-P_{max})\,(1.42) = \epsilon\,(C-P_{max})\,\frac{r_S}{r_R}\left(1+\left(\frac{D_{max}}{h_R}\right)^{k_R}\right), \\ &> \epsilon\,(C-P_{max})\,\frac{r_S}{r_R}\left(\frac{1+\left(\frac{(t-\tau_R)}{h_R}\right)^{k_R}}{1+\left(\frac{D(t-\tau_S)}{h_S}\right)^{k_S}}\right). \qquad (7)\end{aligned}$$

Therefore, if the probability of mutation to resistance is 0.001 then the starting OD for the resistant strain would have to be less than 0.0002 (for $P_{max} = 0.2$) and less than 0.00035 (for $P_{max} = 0.1$) before there is any risk of mutational costs exceeding competitive benefits.

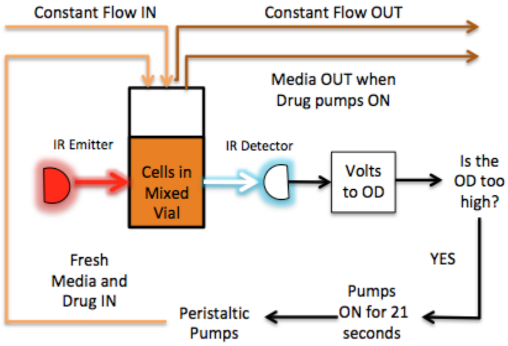


**Fig S1. Computer controlled bioreactors** Constant volume bacterial cultures (17 mL) are grown in glass vials with customized Teflon tops that allow inflow and outflow of fluid via silicone tubing. Flow is managed by a series of computer-controlled peristaltic pumps which are connected to media and drug reservoirs. Cell density is monitored by light scattering using infrared LED/Detector pairs on the side of each vial holder. Voltage readings are converted to optical density (OD) using a calibration curve based on separate readings with a table top OD reader. Up to 9 cultures can be grown simultaneously using a series of multi-position magnetic stirrers. The entire system is controlled by custom Matlab software. Flow chart (above) depicts adaptive drug therapy (lower branches) intended to maintain constant OD by adding drug in response to changes in cell density.

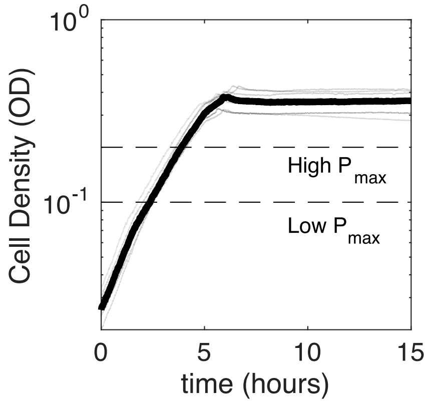


**Fig S2. Growth of resistant cells in unperturbed bioreactors.** Cell density (OD) over time for REL607-derived resistant strains in bioreactors without influx or outflow of media. Transparent black lines correspond do growth curves performed in parallel with each bioreactor experiment. Thick black curve is the median over replicates. Dashed lines indicate threshold densities used in experiments ($P_{max} = 0.2$ and $P_{max} = 0.1$).

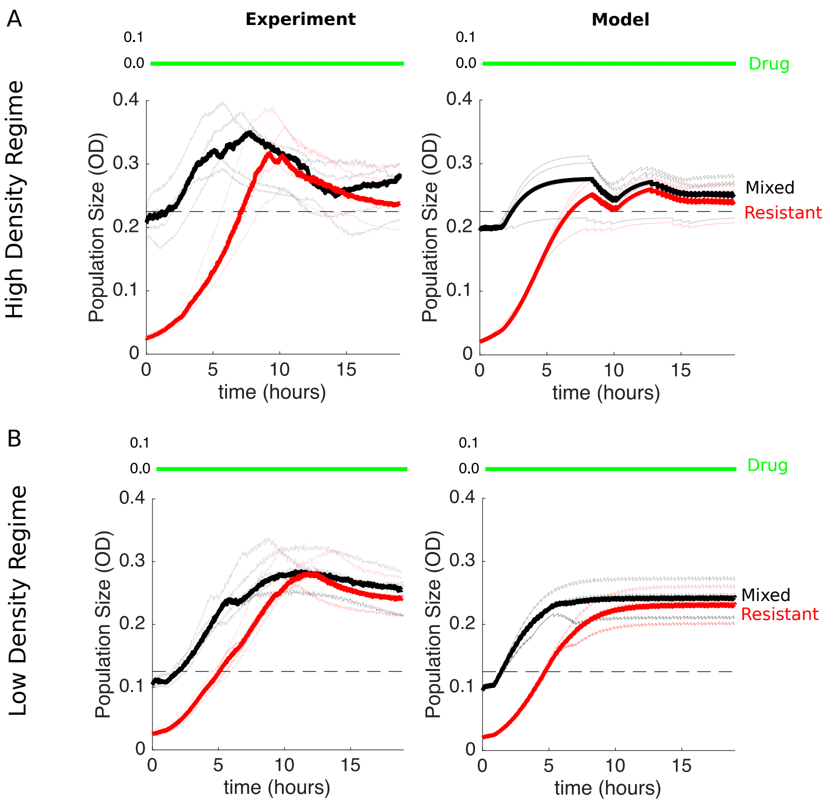


**Fig S3. Matched drug-free control populations are not contained by adaptive dosing protocol**. Conditions are identical to those in Figure 3B and Figure 3C except that all populations receive drug-free media rather than drug solution media as part of the adaptive dosing protocol.

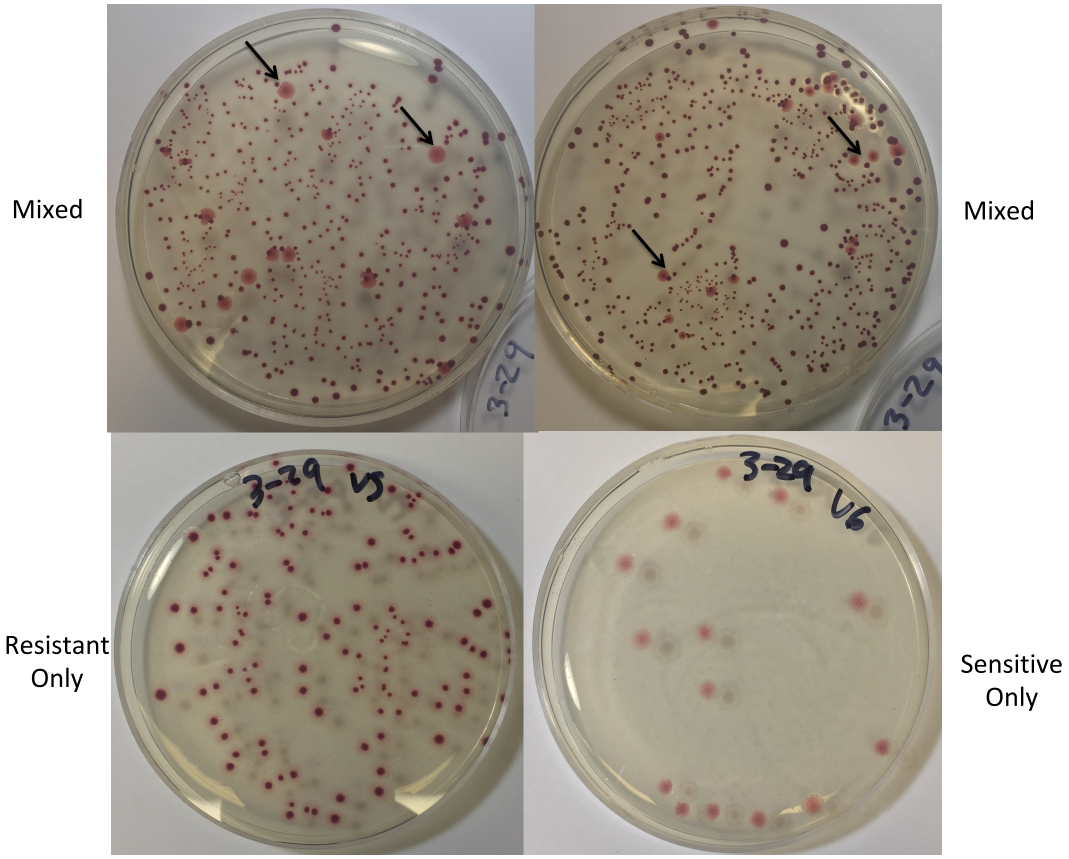


**Fig S4. Mixed populations contain primarily resistant cells at final time point of escape time experiment.** The REL606-derived resistant strain appears red and the sensitive REL607 strain appears pink when grown on tetrazolium arabinose (TA) plates. Upper panels: Samples from two mixed vials taken at the end of a high density escape time experiment (as in Figure 3B). Arrows indicate sensitive colonies. Bottom row: Samples from the end of a high density escape time experiment for a vial seeded with only resistant bacteria (left) and a vial seeded with only sensitive bacteria (right).
